# Agonist and antagonist diverted twisting motions of single TRPV1 channel

**DOI:** 10.1101/2020.08.18.255109

**Authors:** Shoko Fujimura, Kazuhiro Mio, Masahiro Kuramochi, Hiroshi Sekiguchi, Keigo Ikezaki, Muneyo Mio, Kowit Hengphasatporn, Yasuteru Shigeta, Tai Kubo, Yuji C. Sasaki

## Abstract

Transient receptor potential vanilloid type 1 (TRPV1) channels are activated by heat, vanilloids, and extracellular protons. Cryo-EM has revealed various conformations of TRPV1, and these structures suggest an intramolecular twisting motion in response to ligand binding. However, limited experimental data support this observation. Here we analyzed the intramolecular motion of TRPV1 using diffracted X-ray tracking (DXT). DXT analyzes trajectories of Laue spots generated from attached gold nanocrystals, and provides picometer spatial and microsecond time scale information about intramolecular motion. We observed that both an agonist and a competitive antagonist evoked rotating bias in TRPV1, though these biases were in opposing directions. Furthermore, the rotational bias generated by capsaicin was reversed between the wild type and the capsaicin-insensitive Y511A mutant. Our findings bolster the understanding of the mechanisms used activation and modulation of TRP channels, and this knowledge can be exploited for pharmacological usage such as inhibitor design.

## Introduction

Transient receptor potential (TRP) channels constitute a superfamily of nonselective cation channels, including 28 different genes in human. TRPV1 plays a key role in nociception and is activated by various stimuli, including heat, low pH, and vanilloids (*1-3*). Structures of TRPV1 have been determined by cryo-EM at near-atomic resolution (*4, 5*), and they provide a fundamental framework for TRP activation. TRPV1 is composed of a homotetramer, and each subunit includes six transmembrane segments and a pore-forming region. Transmembrane segment 5 (S5), the pore helix, and transmembrane segment 6 (S6) together form a pore in the assembled tetramer, surrounded by a voltage sensor-like (VSL) domain composed by a bundle of four transmembrane helices (S1–S4). Both the N- and C-termini are located inside the cell, and a large ankyrin repeat domain is present at the N-terminus.

Cryo-EM revealed conformational dynamics of TRPV1 induced by capsaicin, capsazepine, resiniferatoxin (RTX) from *Euphorbia resinifera*, and a double knot toxin (DxTx) from Chinese bird spider venom (*4, 6*). Indeed, these cryo-EM data, as well as simulation studies (*7, 8*) suggest rotational and twisting motion may occur during the gating, but direct detection of this movement has remained technically challenging.

## Results

### Single molecule analysis of TRPV1 using DXT

Our studies used full-length human TRPV1 which was subcloned into pcDNA3.1 and tagged with (His)_6_ sequence at the N-terminus, and tagged with the FLAG sequence (DYKDDDDK) at the C-terminus (FLAG-TRPV1) (Fig. S1A). The function of intracellular Ca2+-permeable FLAG-TRPV1 and mutant was assessed by cell-based Ca2+ flux assay in HEK-293 cells (Fig. S1B).

To understand the intramolecular motion of the TRPV1 upon gating, we used a diffracted X-ray tracking (DXT) method (*9*). DXT uses broadband irradiation of X-rays (Fig. S2), and is based on the principle of Bragg’s law (*10, 11*). The diffraction rings from the X-ray beams, also called Debye Schetter rings, broaden according to their energy bandwidths, and the motion of diffraction spots were analyzed in two rotational axis views, including tilting (*θ*) and twisting (*χ*) angles (Fig. 1A-C). The DXT measurement was performed at BL40XU beamline in SPring-8 Japan. Data were recorded with 100 microsecond (μsec) per frame and a total measurement time was 10 millisecond (ms).

**Figure 1.**
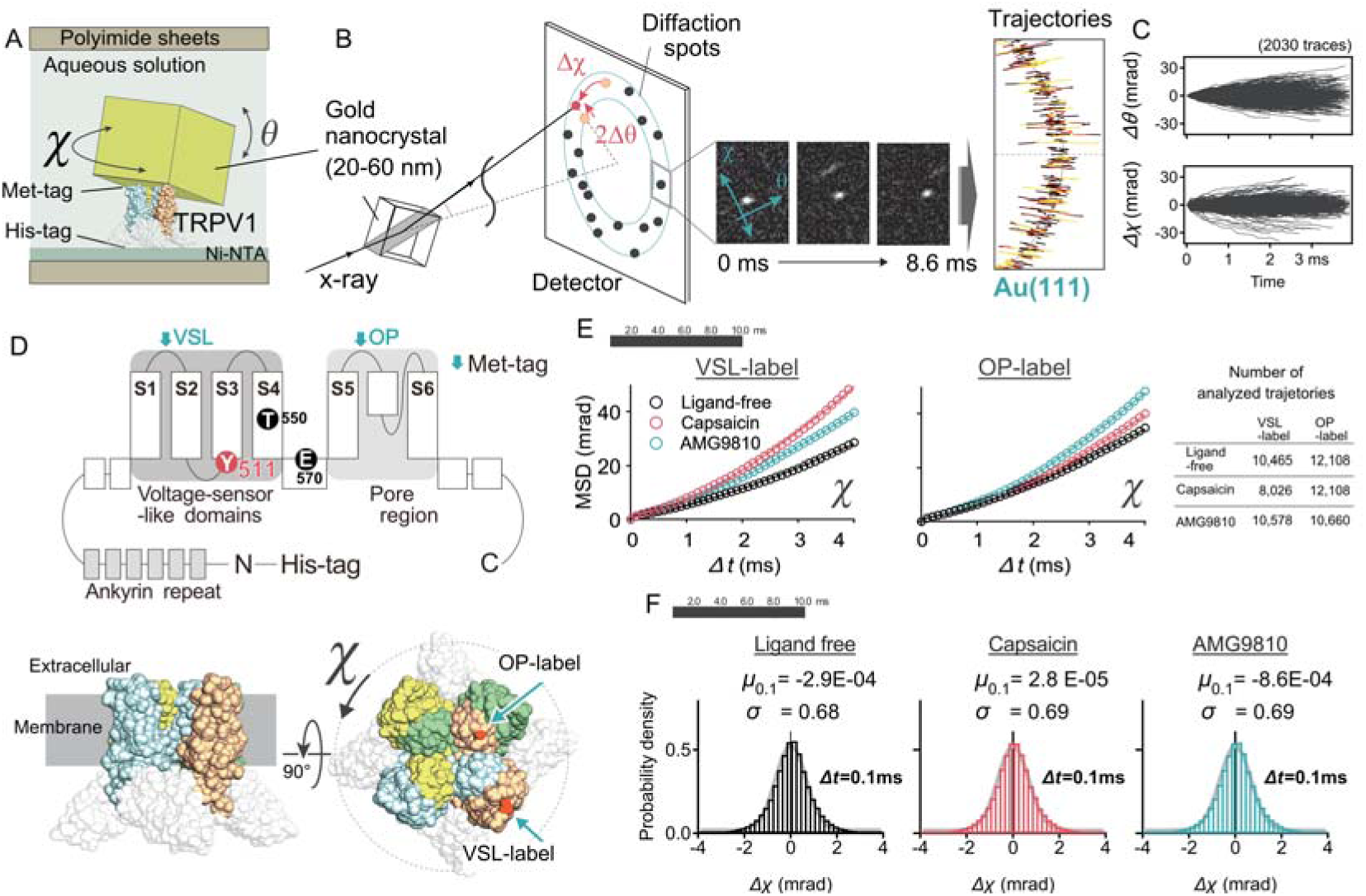
Experimental design for DXT measurement of TRPV1. **(A)** Side view of the sample chamber. N-terminally (His)_6_-tagged TRPV1 was immobilized through a Ni-NTA (self-assembled monolayer formed on the polyimide films. Gold nanocrystals were attached at the extracellular side of TRPV1 via Met-tags introduced in the sequence. **(B)** Schematic of the DXT measurement. Pink-beam X-rays from the SPring-8 synchrotron radiation facility elicited trackable diffraction spots from the gold nanocrystals. The trajectories were projected and analyzed on the *χ-θ* coordinates, separately. The central three panels are representative motion of diffraction spot. The right panel shows piled data of Au (111) spots, representing trajectories. **(C)** Rotational angular displacement along *θ* and *χ* axes over time (4 ms). The 2,030 traces obtained from the ligand-free condition are presented. **(D)** Labeling positions of 2D diagram (upper panel) and 3D structures (lower panels) of TRPV1. The green arrows show Met-tags placed either at the outer pore (OP) region or voltage-sensor-like (VSL) domains of the extracellular side of TRPV1. The gold nanocrystals were specifically bound to the genetically introduced Met-tag (MGGMGGM) in TRPV1 through Au-S binding. Highlighted residues in the 2D diagram are implicated in capsaicin-induced gating properties. **(E)** Mean square displacement (MSD) curves of TRPV1 motion for the *χ* axis. **(F)** Distributions of displacement over an interval time *t* (*Δt* = 0.1 ms) for the VSL-label. Histograms are fitted by the normal distribution (thick gray line), in which the mean *μ* and standard deviation *σ*.

FLAG-TRPV1 was immobilized through a Ni-NTA self-assembled monolayer formed on polyimide films (Fig. 1A). To provide direct visualization of the intramolecular dynamics of TRPV1, the extracellular side of FLAG-TRPV1 was labeled with gold nanocrystals through Au-S binding via Met-tags (MGGMGGM) introduced in the sequence. AFM imaging shows the size of gold nanocrystals were ranging from 40 to 80 nm diameter (*10*). Two extracellular regions were separately labeled with gold nanocrystals: the outer pore (OP) region and the VSL domain (Fig. 1D). The rotation angle (*χ*) of the labeled position was determined relative to the immobilized cytoplasmic domains when viewed from the extracellular side (Fig. 1D, lower).

The time-averaged mean square displacement (MSD) (*12*) curves were obtained for both the VSL- and OP-labeled positions of TRPV1 (Fig. 1E). At the VSL position, capsaicin enhanced the slope of the MSD curve. Interestingly, the slope was also increased by the competitive antagonist AMG9810 (*13*). The diffusion constants (*D*_*χ*_) of capsaicin and AMG9810 obtained from the MSD curves were increased by 46% and 41% over that of the control (ligand-free), respectively (Table S1). The *D*_*χ*_s at the OP-labeled position were also increased by capsaicin and AMG9810, but they were much smaller than those observed at the VSL-label (Table S1) and the increasing rates were limited to 2% and 7% of the control, respectively.

The probability distributions of *χ* were plotted as a function of time-interval (*Δt*) to investigate the dynamics of TRPV1 motion, known as the van Hove self-correlation function (VHC) (*14*). For the ligand-free state, the distribution at *Δt* = 0.1 ms (Fig. 1F) and other time intervals (Fig. S3) fitted well with the Gaussian function. The standard deviation (*σ*) of the Gaussian function was increased from 0.51 to 0.55 by capsaicin (Fig. 1F). The mean *μ* values of the Gaussian function were very small in all conditions (−5.4E-03, 5.3E-05 and 3.6E-03 mrad for ligand-free, capsaicin, and AMG9810, respectively) and there was no bias in the shape of the distribution (Fig. 1F and S4). From the distributions using all detected trajectories, we could not find any significant rotational bias in TRPV1.

### Analysis using lifetime classification

Because the velocities of Brownian particles diverge when time intervals approach zero (*15, 16*), and because the trajectories are not differentiable anywhere due to its discontinuities, a lifetime was introduced to classify the motion of trajectories (Fig. 2A-C, and Supplementary Video1, Video2). An inverse relationship was clearly shown between the lifetime and the average step sizes at 0.1 ms intervals; the longer the lifetime, the smaller the step size (Fig. 2D).

**Figure 2.**
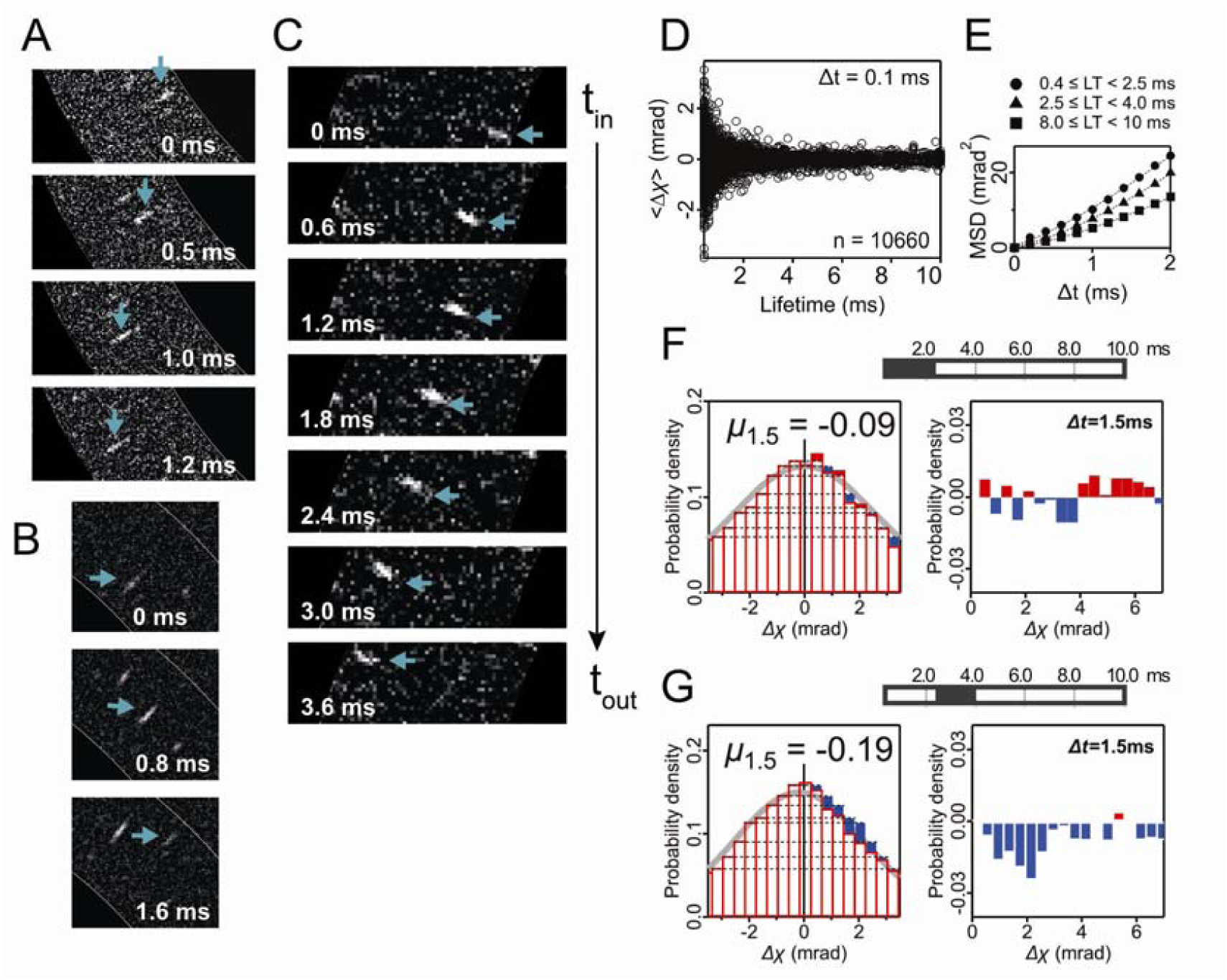
Velocity-based classification. **(A-C)** Trajectory data. **(A, B)** Representative short-lived spots with 1.2- and 1.6-ms lifetime, respectively. **(C)** Representative long-lived spots (blue arrows) with 3.6-ms lifetime. **(D)** Lifetime vs. average step sizes over 0.1-ms time-interval. Presented data of OP-label with capsaicin. **(E)** The MSD of OP-label with capsaicin. The diffusion constants were 9.87, 7.30 and 5.04 mrad^2^ / ms, for short (LT < 2.5 ms), long (2.5 ≤ LT < 4 ms), and the longest (8 ≤ LT < 10 ms) lifetime groups. respectively. **(F, G)** Representative probability density at *Δt* = 1.5 ms (left panels) and subtraction of the probability density (right panels).

We applied statistical analysis into three groups: short [lifetime (LT) < 2.5 ms], long (2.5 ≤ LT < 4 ms), and the longest (8 ≤ LT < 10 ms) lifetime groups. The diffusion constants *D*_*χ*_ of OP-label with capsaicin were obtained from the MSD curves; 9.87, 7.30 and 5.04 mrad^2^ / ms, for short, long, and the longest group, respectively (Fig. 2E and Table S2). In the short lifetime trajectories, we could not find any statistical significance in the mean *μ* values among the experimental conditions (Fig. 2F, upper and Fig. S5). However, the mean *μ* value of OP-label was decreased to –0.19 by capsaicin in the long lifetime trajectories (Fig. 2G). Subtraction of the probability density showed clearer rotational bias in the long lifetime group than the short lifetime group (Fig. 2F and 2G).

The mean *μ* values plotted over time-interval *Δt* (denoted as mean plot) represent the rotational bias generated in TRPV1 (Fig. 3). Rotation angle was determined from the extracellular side, by the motion of the extracellular OP- and VSL-region in contrast to the cytoplasmic domain, which was immobilized to polyimide film. The motions of the short lifetime trajectories were again shown to be non-biased (Fig. 3A and Fig. S5). In contrast, the long lifetime trajectories showed unique rotational biases to their sample conditions (Fig. 3B). We judged the biased motion to be clockwise (CW), counterclockwise (CCW), and non-directed movement from the mean plots. From previous studies, a CW bias is considered to be a motion toward channel opening (*4*). The maximum rotation angles of capsaicin-induced TRPV1 were 1.10 and 1.55 degrees at 1 ms-intervals to the CW direction both at the OP and VSL positions, respectively (Fig. 3C).

**Figure 3.**
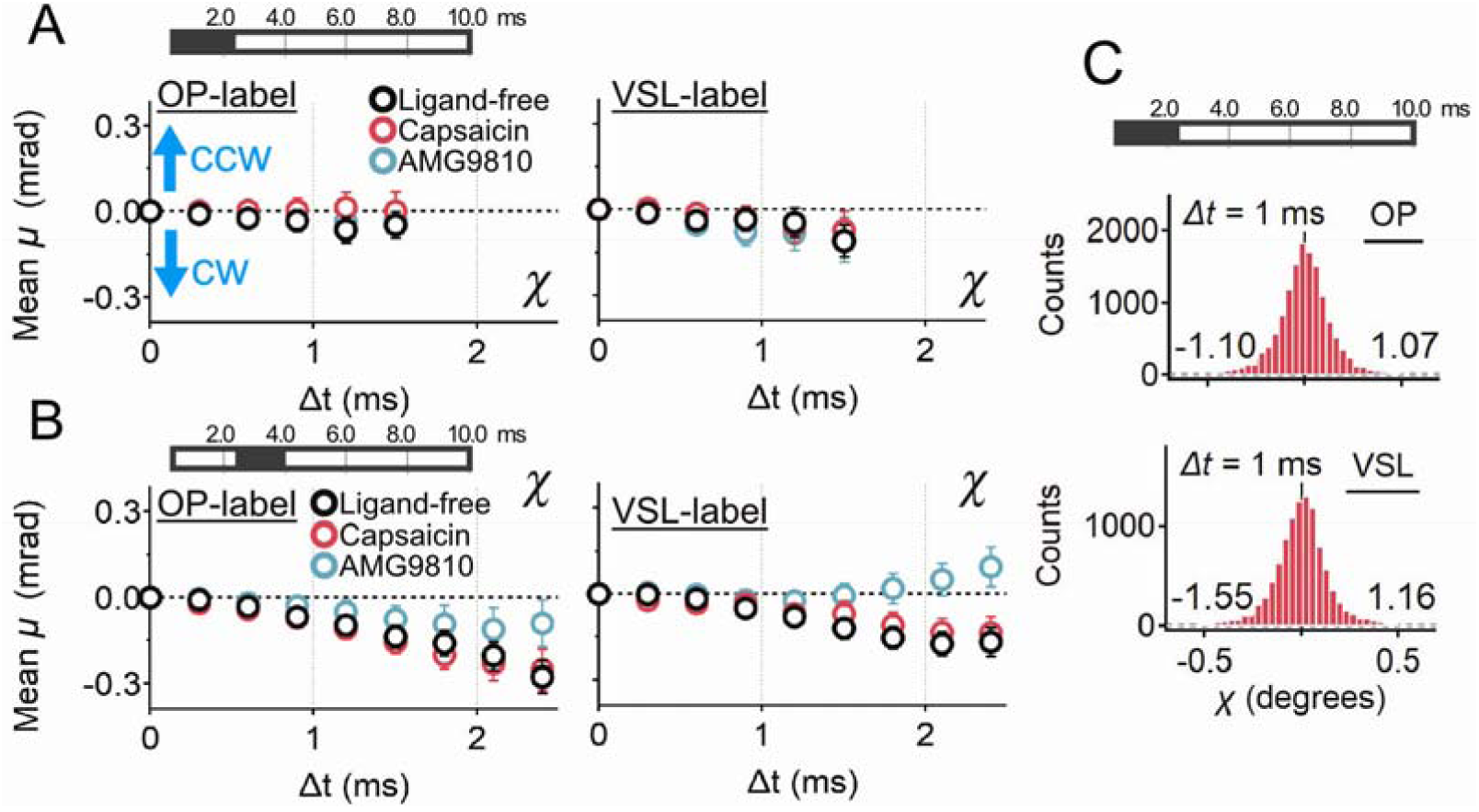
Mean plot of wild type TRPV1. **(A)** The mean plot and 95% confidence intervals for the angle *χ* at short lifetime group (LT < 2.5 ms, left) The rotation angle was mostly non-directed or slightly CW biased at both the OP-label (left) and VSL-label (right). **(B)** The mean plot at long lifetime group (2.5 ≤ LT < 4 ms) of OP-label (left) and VSL-label (right). Dark-filled periods of the top bars show the lifetime to which the data belong. **(C)** The distribution of angular displacement in 1 ms time interval. Data are displayed for capsaicin. Maximum and minimum values of angular displacement are shown at lower right (CCW bias) and left (CW bias) of each plot, respectively.

At the OP-label, all experimental conditions showed CW rotational bias. However, the mean *μ* value at *Δt* = 2.4 ms (*μ*_2.4_) of ligand-free and capsaicin were equivalent (*μ*_2.4_ = −0.23 and −0.25, respectively), while *μ*_2.4_ of AMG9810 was much smaller (*μ*_2.4_ = −0.09). As for the VSL-label, ligand-free and capsaicin binding also showed CW rotational bias, while AMG9810 showed bias in the CCW direction (*μ*_2.4_ = 0.10).

### Motion analysis of the slowest-moving group

We also analyzed the movement of the slowest moving group using the longest life filtering (8 ≤ LT ≤ 10 ms). Capsaicin-induced movement clearly showed the CW bias, while the movement of the ligand-free condition was CCW biased at both the OP and VSL positions (Fig. 4). This result shows that the capsaicin evoked relatively slow movement in TRPV1, which may persist up to 10 ms.

**Figure 4.**
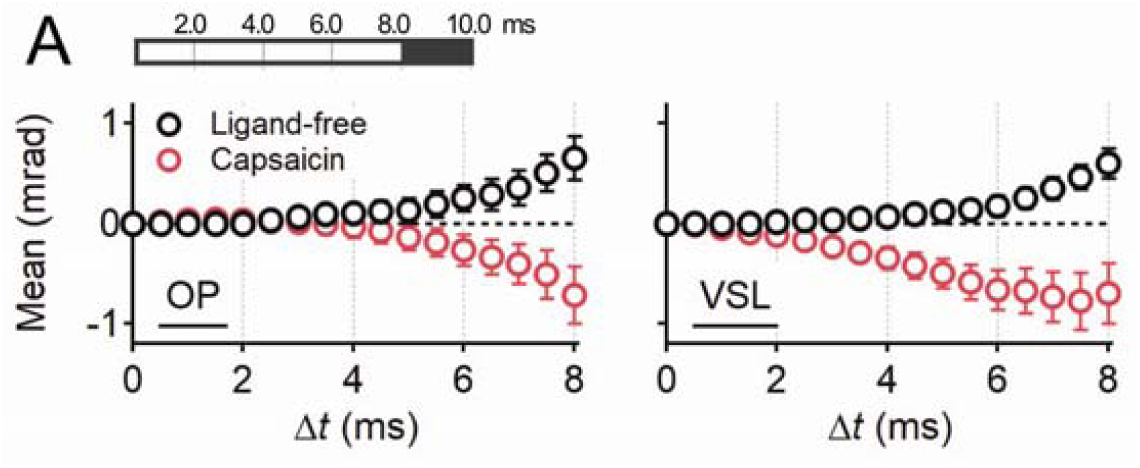
The rotational motion of the slowest-moving group. The rotational motion of ligand-free and capsaicin-bound TRPV1 in the longest lifetime-filtering (8 ≤ LT < 10 ms). Capsaicin showed CW, while ligand-free showed CCW biased at both the OP-label (left) and VSL-label (right). The mean plots (circles) and 95% confidence intervals (bars) of the 8-10 ms lifetime group were presented. The number of trajectories used for the analysis were: for 371 (OP of ligand-free), 296 (OP of capsaicin), 353 (VSL of ligand-free) and 177 (OP of capsaicin).

### Motion analysis of capsaicin-insensitive mutant

To confirm whether the CW bias was really associated with channel gating, we examined the motion of the capsaicin-insensitive Y511A mutant. Trajectories of gold nanocrystals at both OP- and VSL-label were also obtained from the Y511A mutant, and were similarly classified into short (LT < 2.5 ms) and long (2.5 ≤ LT < 4 ms) lifetime groups. As was the case with the wild type channel, the short lifetime groups did not show any specific rotational bias (Fig. S6). As for the long lifetime groups, the rotational movements of Y511A in the ligand-free condition were non-biased (OP-label) or slightly biased toward CW (VSL-label) (upper panels of Fig. 5A). However, capsaicin evoked a strong bias toward the CCW direction in Y511A both at the OP- and VSL-label positions (lower panels of Fig. 5A). This was a big difference compared to the capsaicin-induced CW bias of the wild type. Capsaicin-induced CCW bias in the Y511A mutant resembles AMG9810-induced CCW bias in the wild type channel.

**Figure 5.**
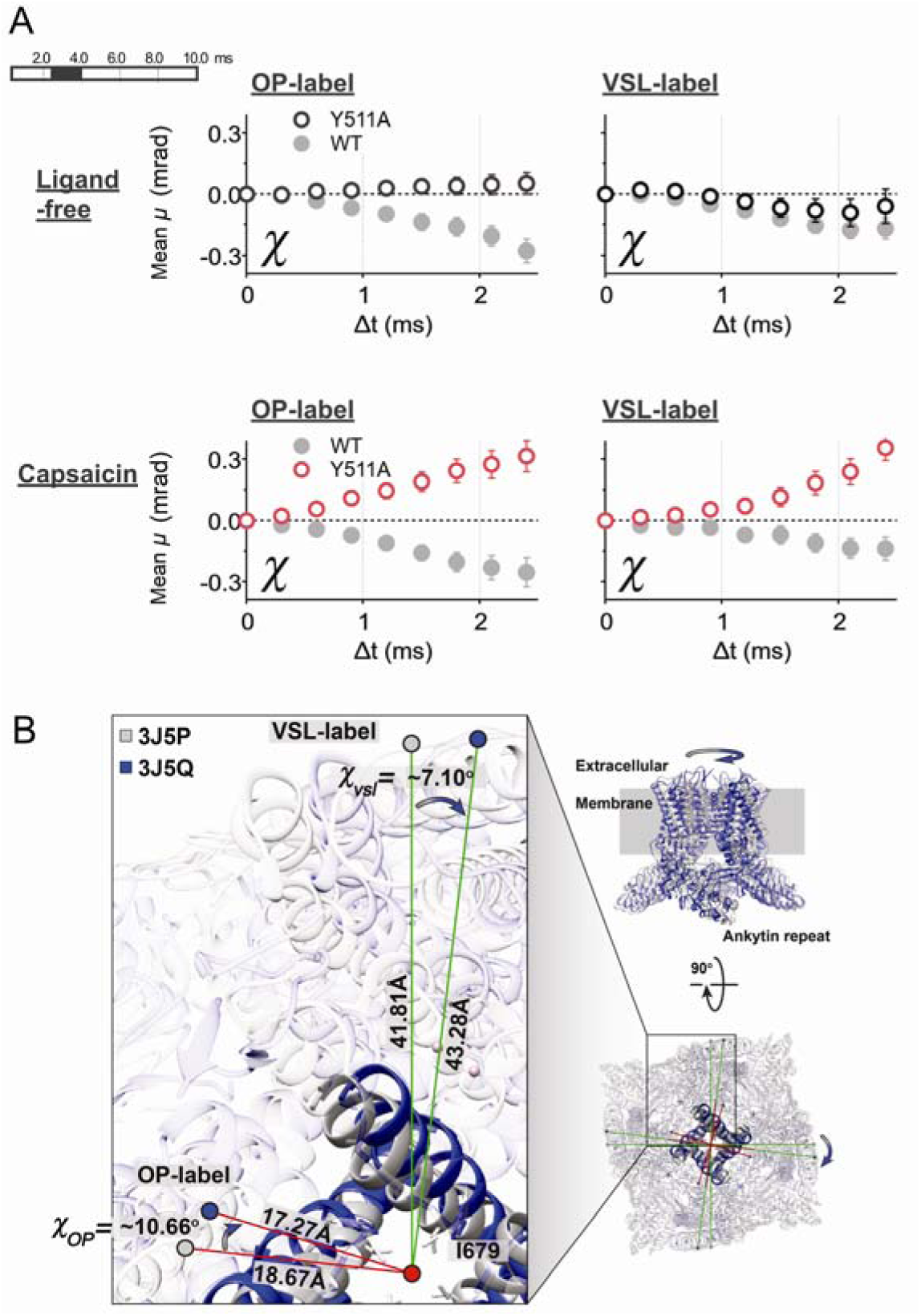
Mean plot of the Y511A mutant and structural comparison of closed and open form of TRPV1. **(A)** The mean plots (circles) and 95% confidence intervals (bars) for the angle *χ* of the long lifetime filtering (2.5 ≤ LT < 4 ms) for the OP-label (left panels) and VSL-label (right panels). Upper panels demonstrate the non-biased motion of Y511A at the ligand-free state (open circles). Lower panels show the capsaicin-induced biased motion of Y511A toward positive (CCW) direction (red open circles). **(B)** Superimposition of apo-TRPV1 (PDB: 3J5P, grey) and DkTx/RTX-TRPV1 (3J5Q, blue). The ankyrin repeat domains were held as a fulcrum. Rotational angles after DkTx/RTX binding was calculated at the glycine 602 (OP-label) and the tyrosine 463(VSL-label). Both the OP- and VSL-positions rotated clockwise direction, resulting in 10.6 and 7.1 degrees, respectively.

## Discussion

It is technically challenging to directly detect twisting motion of channels, because the motion involves rapid conformational changes ranging from the microsecond to millisecond time scale. They are beyond the temporal and spatial resolution of most imaging-based techniques. DXT provide sub-angstrom scale information of conformational dynamics as well as functional intermediates. In this study, a diffusion constant of the glycine 602 at the OP-labeled position, which locates 17.3 Å distance from the center of the pore, was calculated to be 12.8 pm^2^/ms (Fig. 5B).

Because all measurements were performed in the closed sample chambers under the open-closed equilibrium of channel, we could not detect any significant differences in the rotational bias at the first analysis. By applying lifetime filtering, however, we were able to extract the motions related to pore gating. The most prominent difference was observed by the filter 2.5–4.0 ms (Figs. 3A and 3B), where control and capsaicin showed equivalent levels of CW rotation bias, whereas AMG9810 was CCW-biased.

We compared cryo-EM structures of TRPV1 reported in the Protein Data Bank (https://www.rcsb.org). Rotation angle was determined by the movement of glycine 602 for OP-label and tyrosine 463 for VSL-label, when the ankyrin repeat domains was held as a fulcrum. In the comparison of apo- (PDB: 3J5P) and capsaicin-bound TRPV1 (PDB: 3J5R), we and others (*17*) found no obvious conformational change both at the OP- and VSL-positions examined in this experiment (Fig. S7).

We further compared the structures of apo- (PDB: 3J5P) and the DkTx / RTX bound TRPV1 (PDB: 3J5Q) to understand the twisting direction of helices in response to the channel opening. The rotational motion generated by the DkTx / RTX was CW direction both at the OP- and VSL-positions, with calculated angles of 10.6 and 7.1 degrees, respectively (Fig. 5B). TRPV2 (Ref Sec ID: NP_057197) and TRPV3 (NP_001245134) also rotate CW direction in response to ligand binding (*18, 19*) which share 46% identity and 41% similarly in amino acid sequence to TRPV1, respectively. Because the overall architectures of these channels are quite similar to each other, intramolecular motions at channel opening may share similar mechanisms. In considering the lack of obvious structural change at the OP- and VSL-positions by capsaicin, capsaicin-induced rotational biases at these positions must be much smaller than those by DkTx / RTX. Indeed, the capsaicin-induced absolute value of maximum twisting angle determined by DXT was 1.55 degree at Δt = 1 ms (Fig. 3C).

This rotational motion associated with gating may reflects several intermediate states of the channel. Cryo-EM presents three structures of TRPV1 so far; ‘closed‘, ‘partially open‘, and ‘full open’ states. At least two transition states exist along with the pore opening pathway during activation of capsaicin (*17*). In the first pathway, the main transition occurs from the closed state to the partially open state, by capsaicin binding. After that, the second pathway for the channel opening occurs. For the motion of the longest lifetime filtering group (8 ≤ LT ≤ 10 ms), the difference in the mean-plot curves was obvious between the control and capsaicin (Fig. 4). Capsaicin-induced movement was found to be CW, on the other hand, ligand-free was CCW biased. This delayed movement may reflect a capsaicin-evoked second pathway from the partially-open to the fully open state.

AMG9810, a competitive antagonist of TRPV1, interacts with the same binding pocket as capsaicin. Thus, a fundamental question remains why only the agonist causes channel opening. Indeed, AMG9810 was shown to induce CCW rotational bias of TRPV1 in this study. Cryo-EM-based 3D structures and the computation studies (*4, 7, 17*) have suggested how capsaicin activates TRPV1 at the atomic level. Briefly, residues Y511, T550, and E570 mainly contribute to capsaicin binding (Fig. 5B and 1D). Upon binding, capsaicin is stabilized by T550 through Van der Waals interactions, followed by a series of structural rearrangements generating an outward pushing force to the S4-S5 linker by E570 through hydrogen bonding, and opening the S6 activation gate by flipping the aromatic side chain of Y511 (*7*). Loss of the hydroxyl group from the vanillyl group in AMG9810 may abrogate the ability to interact with the S4-S5 linker, while interactions with the binding pocket remains (*7, 20*)

In this study, we found that even the ligand-free state of TRPV1 has a bias toward CW (pore opening) direction, which was not seen in the Y511A mutant proteins (Fig. 3A-B and 5A). Considering the proposed mechanisms, a flip-flop fluctuation of the side chain in Y511 may trigger the rotation movement. Indeed, this flipping motion was demonstrated by MD simulation analysis in the ligand-free form (*21*).

Capsaicin evoked strong CCW (pore closing) bias to Y511A (Fig. 5A), which may be in correlation with the results of AMG9810-induced CCW rotation bias in wild type. A loss of interaction between Y511 and the S6 segment in the Y511A mutant proteins may deflect the rotation bias to the CCW direction.

Determining of structural changes at single-molecule level, together with the single channel patch clamp techniques, could be expected to elucidate the exact role of channels such as allosteric mechanism and desensitization of TRPV1.

## Materials and Methods

### Plasmid Construct and protein purification

Full-length human TRPV1 was subcloned into pcDNA3.1 and tagged with (His)_6_ sequence at the N-terminus, and tagged with the FLAG sequence (DYKDDDDK) at the C-terminus (Fig. S2A). Met-tag (MGGMGGM)was introduced at an extracellular loop between the S1 and S2 segments (for VSL-labeling), or a loop between the S5 segment and the pore-forming domain (for OP-labeling). The TRPV1 protein was expressed in HEK293F cells, and purified with two step purification by an anti-FLAG M2 affinity chromatography (Sigma) and a Superdex 200 size exclusion chromatography (GE Healthcare, IL). The peak fractions were concentrated to 0.1 mg/mL for DXT (Fig. S2C). Alanine substitution at the 511 tyrosine was made by site-directed mutagenesis using the QuikChange mutagenesis kit (Agilent Technologies).

### Sample preparation for DXT assays

Gold nanocrystals were obtained via epitaxial growth on a KCl (100) surface and dissolved by a buffer containing 50 mM MOPS (pH 7.0) and 5 mM n-decyl-β-D-maltoside (Dojindo, Japan). The basement surface of the polyimide film (Du Pont-Toray, Japan) was coated with Ni-NTA self-assembled monolayer using dithiobis (C2-NTA) (Dojindo). The 20 *μ*L (His)_6_-tagged TRPV1 (∼0.1 mg/mL) was immobilized on the 50-*μ*m-thick polyimide film. After incubation for 6 h at 4 °C, excess protein was washed out with buffer. The gold nanocrystals, which specifically bind to the genetically introduced Met-tag (MGGMGGM) via Au-S binding, were applied. After incubation for 60 min, excess gold nanocrystals were washed out with buffer. After applying 10 *μ*L of phosphate buffered saline containing 10 *μ*M capsaicin or 10 *μ*M AMG9810, the sample chamber was covered with another layer of polyimide film, and sandwiched by stainless steel frames and screw-clamped. Capsaicin and AMG9810 were purchased from Sigma.

### DXT measurements

X-rays from the beamline (BL40XU, SPring-8, Japan) with energy widths ranging from 14.0–16.5 keV were used for DXT measurements (Fig. S1). The beam size was adjusted to 50[*μ*m in diameter by inserting a pinhole aperture upstream of the sample, and time-resolved diffraction images from gold nanoparticles immobilized on TRPV1 were recorded by an X-ray image intensifier (V7339P, Hamamatsu photonics) and CMOS camera (Phantom v2511, Vision Research). Data were recorded at 100 μsec/frame intervals, and measured 100 frames per spot. For each sample, diffractions at 72 spots were recorded. The distance between the sample and a detector was set to 60 mm. All DXT measurements were carried out at room temperature.

### Data Analysis

The trajectories of diffraction spots on the image plane were tracked by TrackPy (v0.3.2 https://doi.org/10.5281/zenodo.60550) package with python. The trajectories of the diffraction spots were analyzed using custom software written within IGOR Pro (Wavemetrics, Lake Oswego, OR). Data were recorded with 100 microsecond (μsec) per frame and a total measurement time was 10 millisecond (ms). Data with short-lived (0.1 ≤ lifetime ≤ 0.3 ms) were excluded for reducing the analytical noise, because the number of short-lived data were 2.5 times of data with long-lived (0.4 ≤ lifetime ≤ 10 ms).

## Acknowledgments

This work was supported by JSPS KAKENHI Grant Numbers JP17H05539, JP18K06601, JP26102748 and by JST CREST Grant Number JP18071859, Japan. DXT and DXB experiments were performed with the approval of the Japan Synchrotron Radiation Research Institute (Proposal Nos 2017A1140, 2018A1417 and 2019A1498), and of the Photon Factory Program Advisory Committee (Proposal No. 2017G556).

## Supporting information

**Figure S1.**
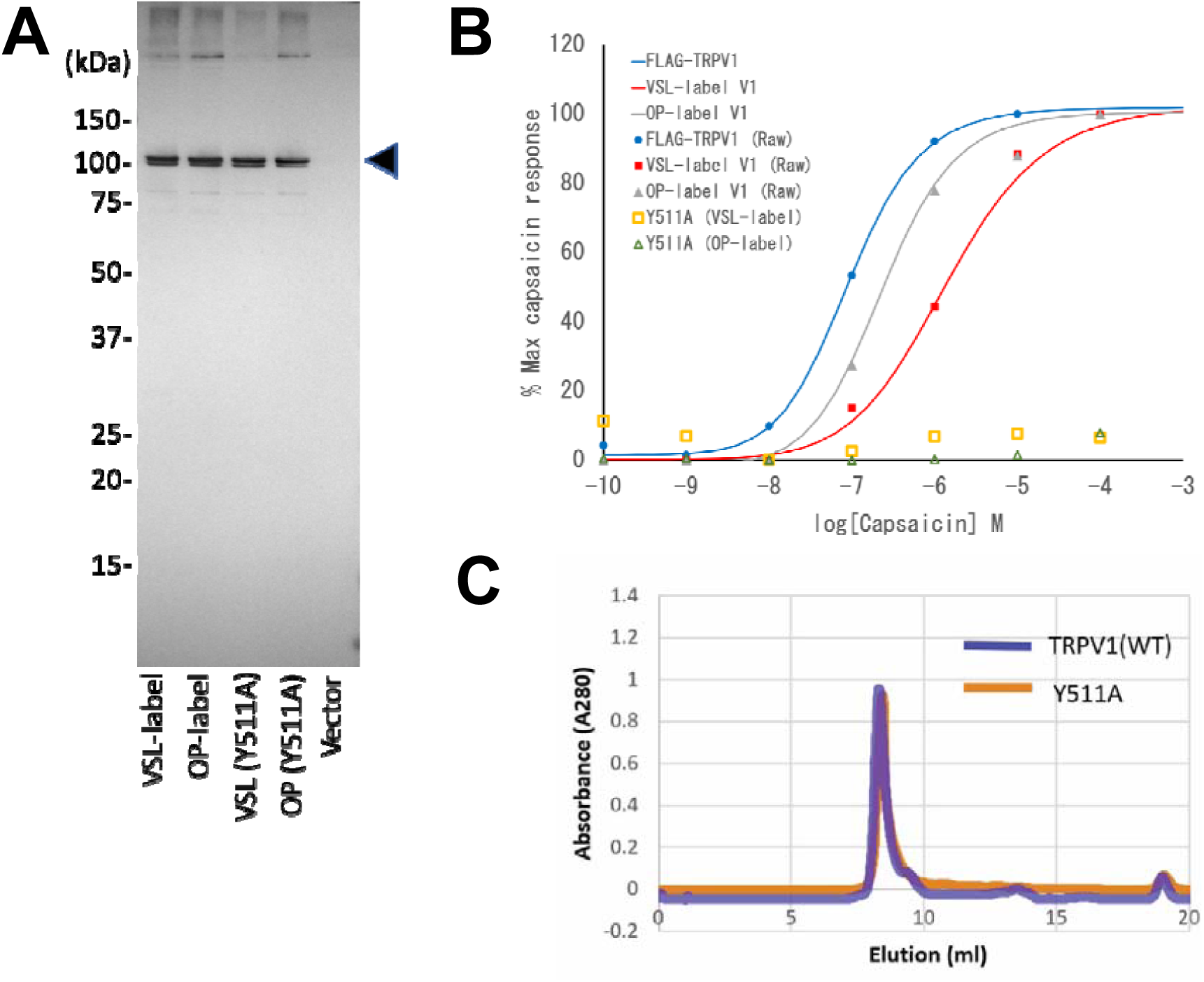
Expression and function of TRPV1 receptors used in this experiment. **(A)** Expression of VSL-label TRPV1, OP-label TRPV1, VSL-label Y511A mutant, and OP-label Y511A mutant on the HEK293 cells were confirmed by the Western blotting of the anti-FLAG antibodies. The right-end lane is a control transfection by the vector only. The arrowhead shows the position of TRPV1 receptors. (B) Responses to the capsaicin were tested by the calcium influx assay using the FlexStation 3 system with the FLIPR Calcium 6 reagent (Molecular Devices Inc.). Mean values of 2-4 independent measurements were plotted. EC_50_ was 0.088 *μ*M(FLAG-TRPV1), 1.32 *μ*M (VSL-label TRPV1) and 0.245 *μ*M (OP-label TRPV1), respectively. **(C)** Gel-filtration profiles (Superdex 200 column) of TRPV1 purification. Buffer composition was; 20 mM Tris-HCl, pH 7.4, 150 mM NaCl, 50 mM MgCl_2_, 1 mM n-Decyl-β-D-maltoside).

**Figure S2.**
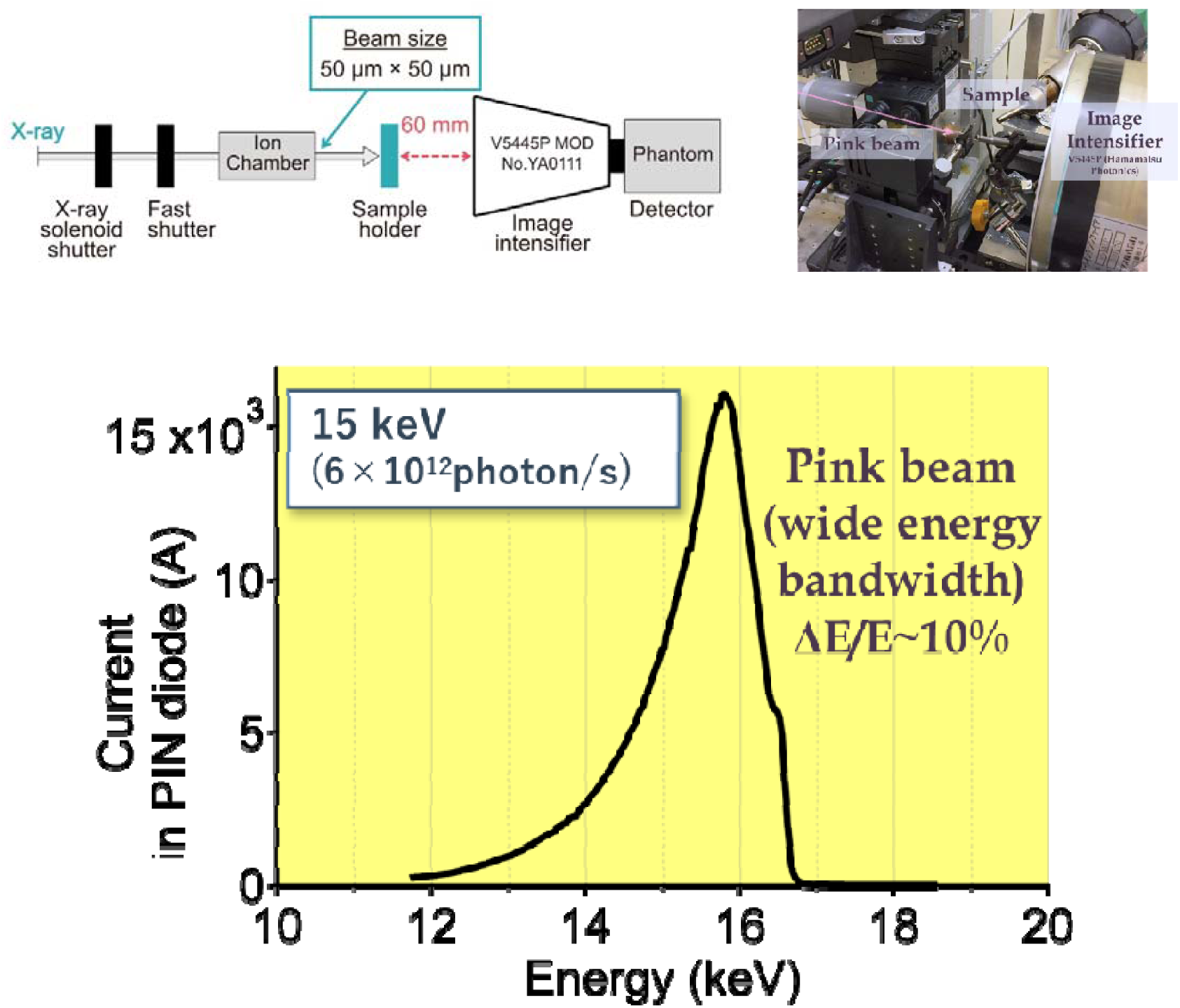
DXT measurement system (upper panels) and the X-ray flux in PIN diode detector for the incident beam (lower panel) X-rays from the beamline (BL40XU, SPring-8, Japan) with energy widths from 14.0-16.5 keV (undulator gap = 30.1 mm) were used for DXT measurements.

**Figure S3.**
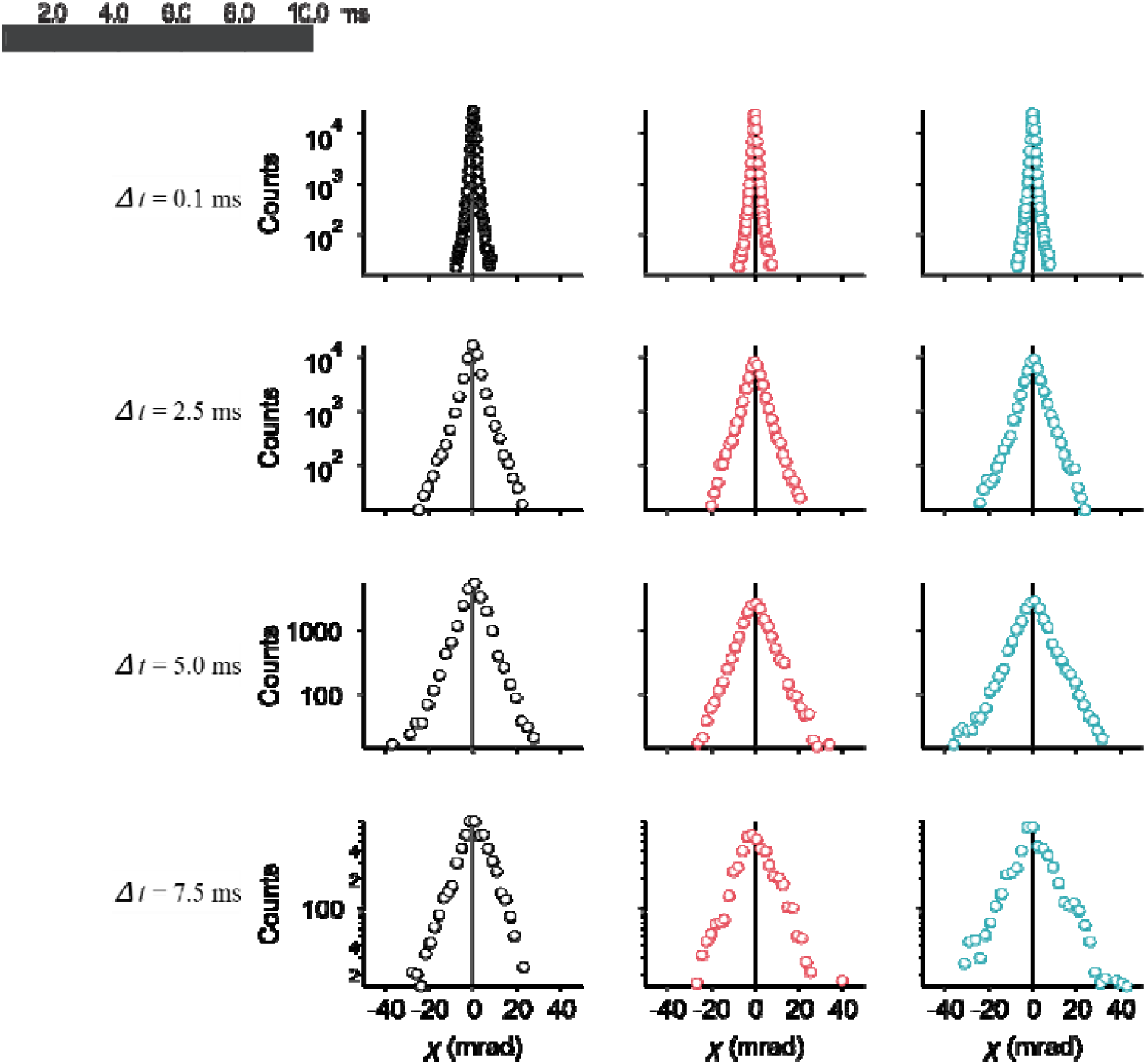
The VHC provide symmetric distribution for OP-label. Before classification of lifetime, the distribution were agreed well by Gaussian function at short time intervals (= 0.1 ms) to large time intervals (= 7.5 ms).

**Figure S4.**
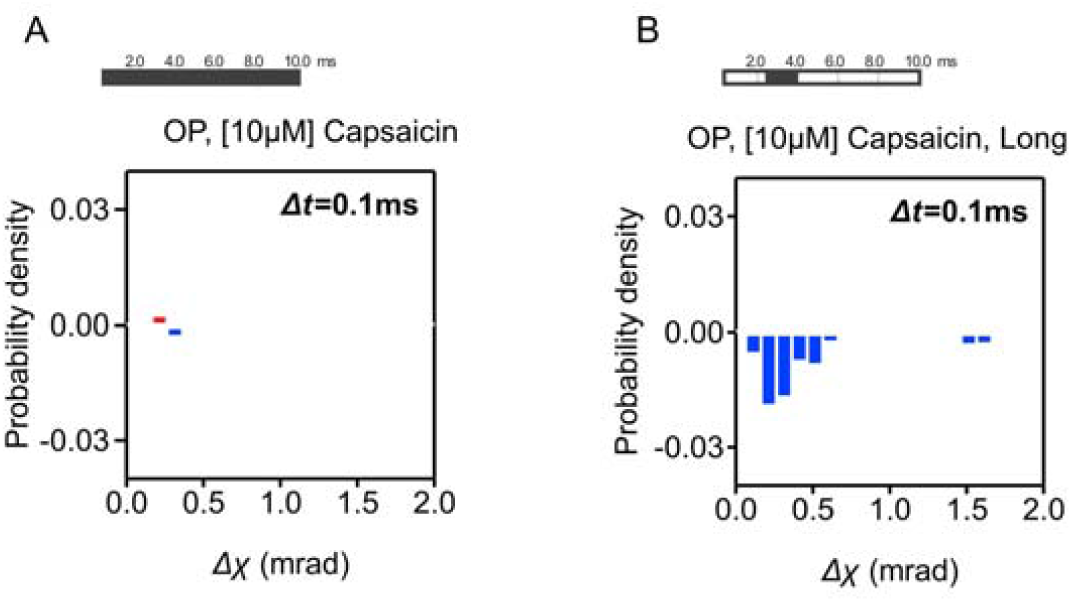
Subtraction in the probability density between the positive value and the negative value for OP-label at *Δt* = 0.1 ms. **(A)** Subtraction of the probability density for all detected trajectories group (0.4 ≤ lifetime (LT) < 10 ms) with capsaicin. **(B)** Subtraction of the probability density for long (2.5 ≤ LT < 4 ms) with capsaicin. Comparing with long LT group which shows significant negative difference, all detected trajectories group didn’t show any obvious rotational bias.

**Figure S5.**
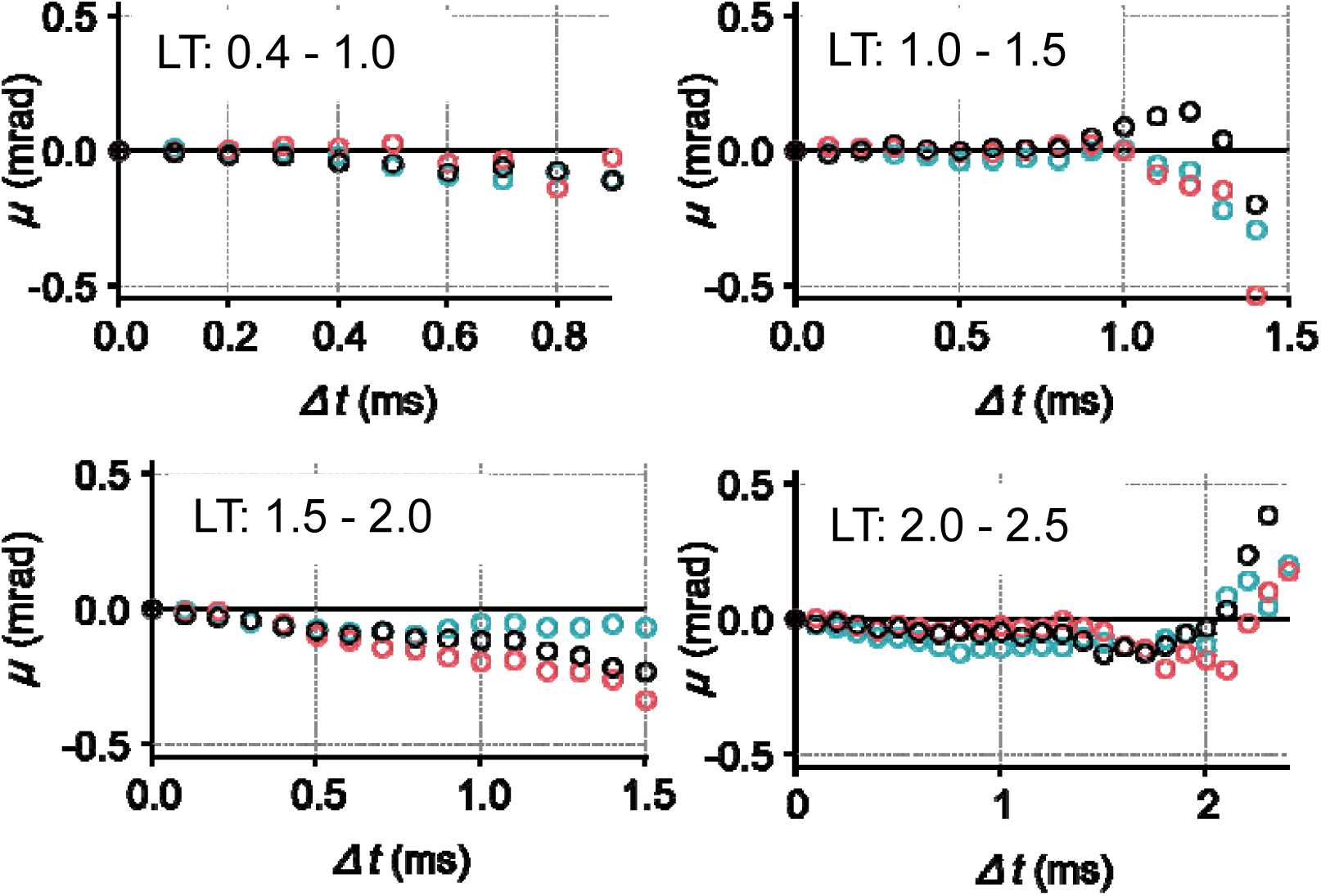
Mean-plot of the VSL-label for the further classified short (the lifetime (LT) < 2.5 ms) lifetime group. Different length of lifetime filtering was further applied to the short lifetime group (LT < 2.5 ms), and the mean-plots of subgroups were calculated. Rotational motion of the subgroups was also confirmed to be non-biased as shown in the result from the short lifetime group (Fig. 3A). Applied lifetime filtering is shown at upper left of each panel.

**Figure S6.**
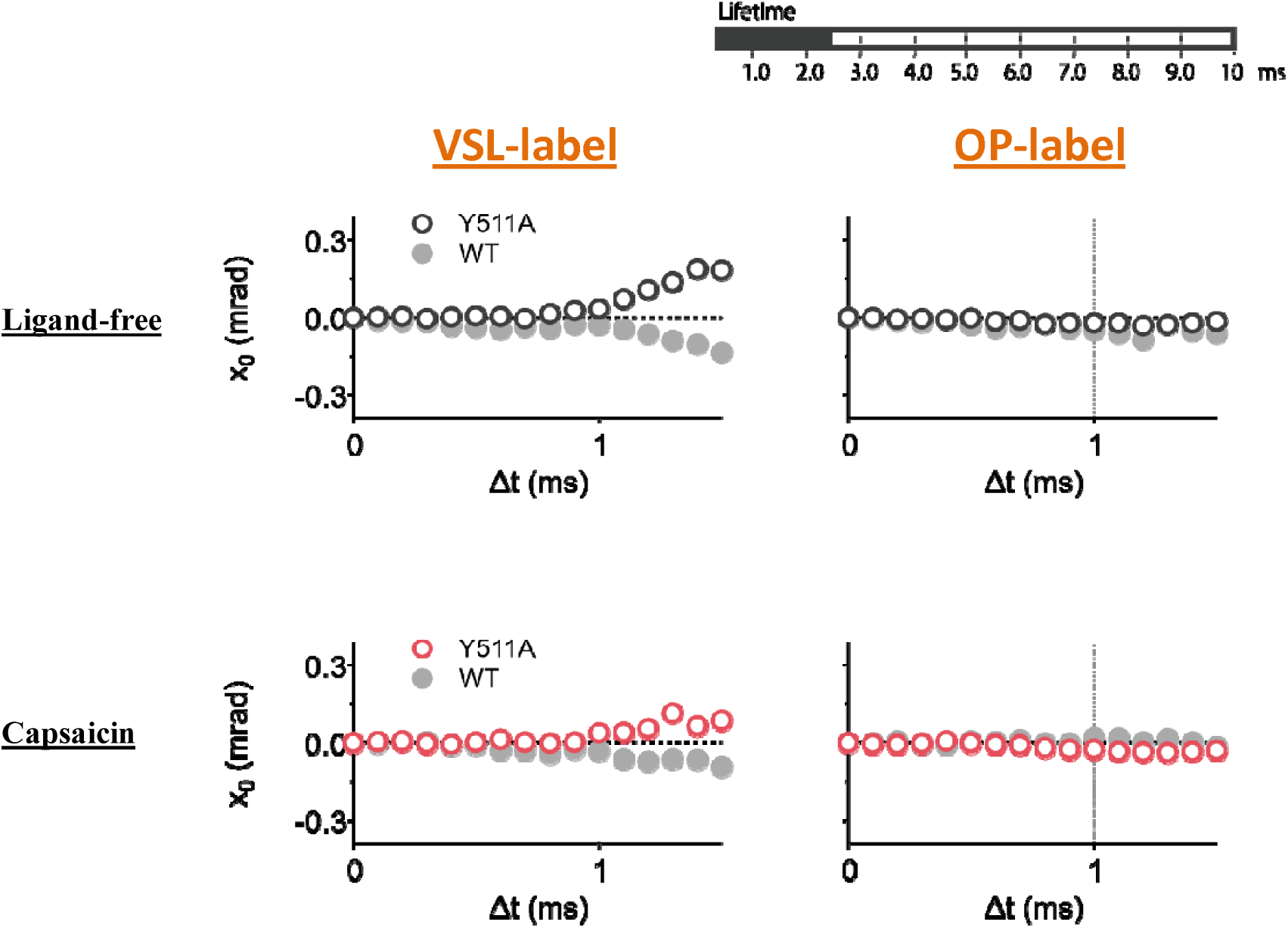
The mean plots of the WT and Y511A against capsaicin for the short lifetime group (LT < 2.0 ms). Like the motion observed in the WT (Fig. 3A), Y511A in the short lifetime groups also did not show any specific rotational bias. Dark-filled area of the top right bar shows the applied lifetime periods.

**Figure S7.**
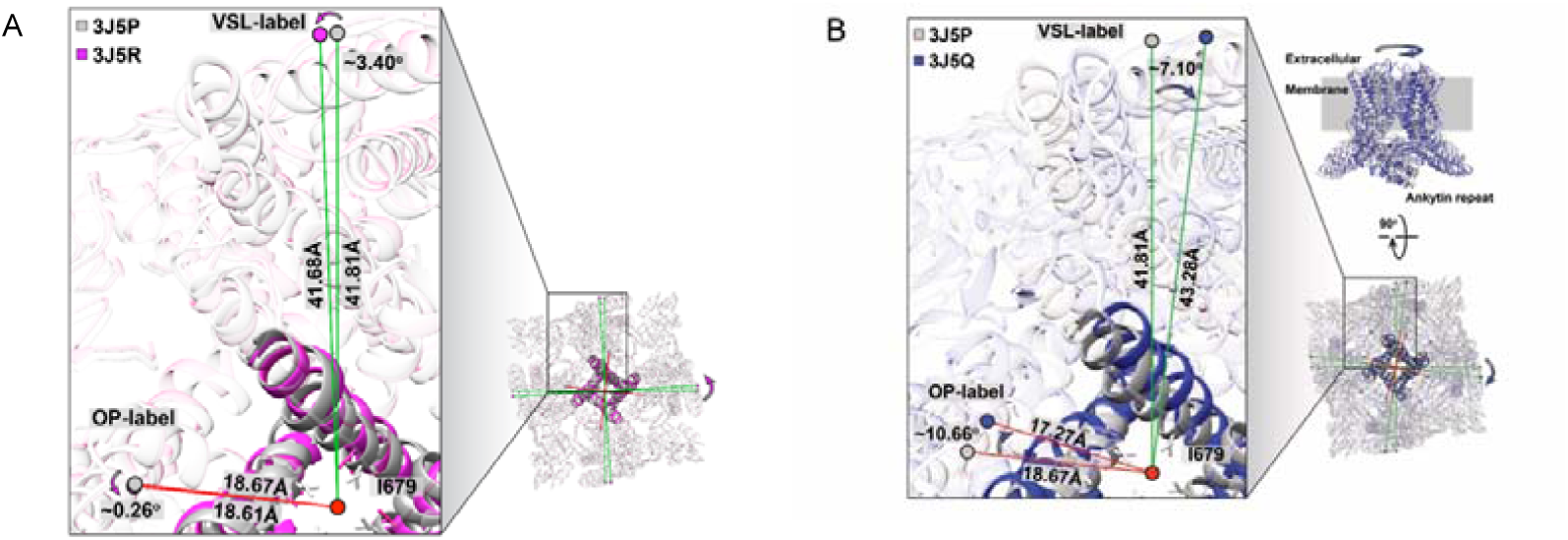
Superimposed views of closed and open-form of TRPV1. (A) The ankyrin repeats domain at residues 111-143 was held as a fulcrum, and the structures were viewed and compared from the extracellular side. Apo (PDB: 3J5P, gray) and capsaicin-bound partially open (3J5R, magenta) structures were superimposed. (B) Apo- (PDB: 3J5P, grey) and DkTx/RTX-bound full-open (3J5Q, blue) structures were superimposed (reposted from the Fig. 5B). No significant conformational change was shown both at the OP- and VSL-positions by capsaicin, while the DkTx/RTX, which induces full-open structure, generated the CW rotation biases. Positions of OP (Gly 602)- and VSL (Tyr 463)-labels were indicated by circles.

**Table S1.**
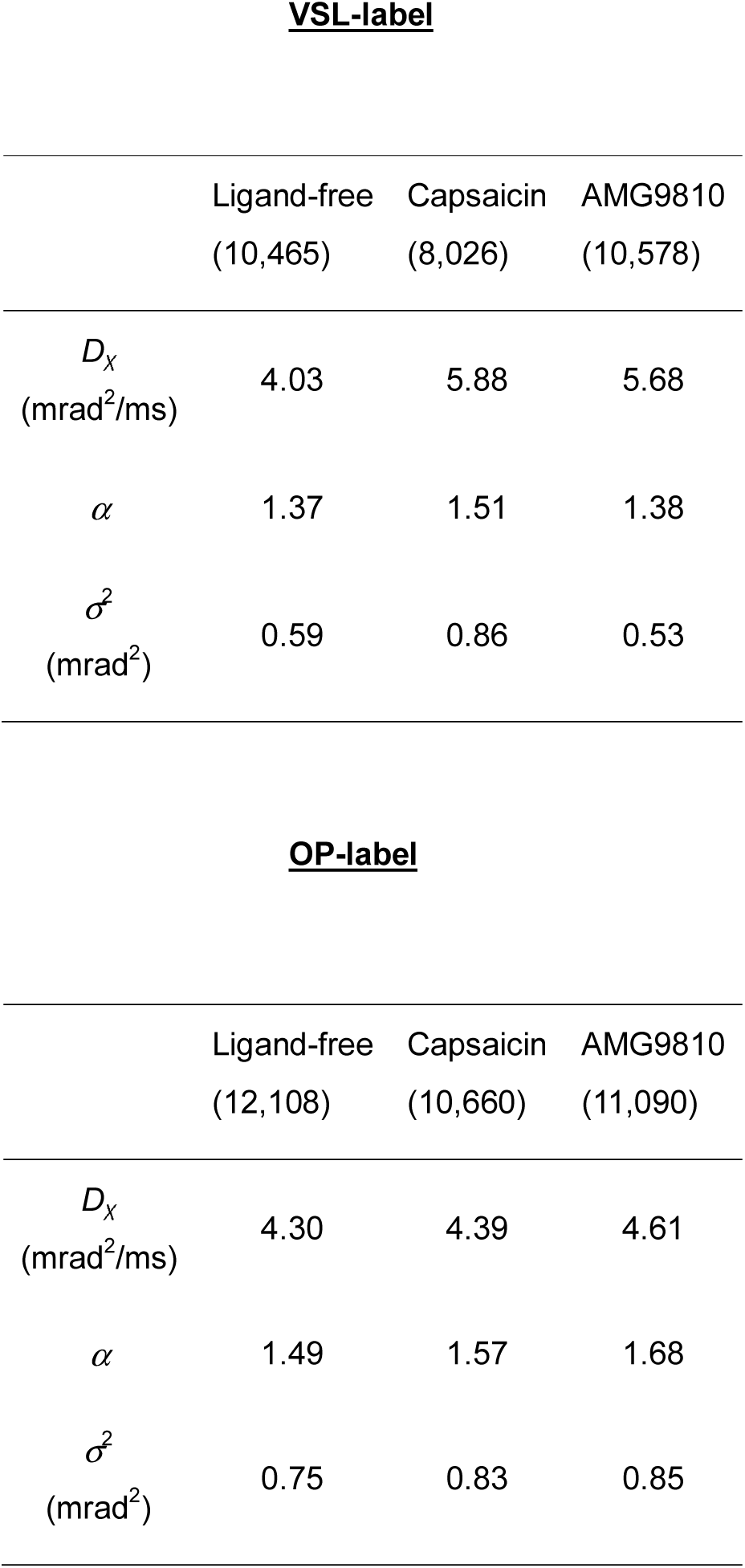
Parameter of MSD for wild type of TRPV1. The MSD curve were fitted by the following function:*δ* ^2^(*t*) = *Dα tα* + 2*β* ^2^. *D*_*α*_ is the anomalous diffusion constant, a non-linear relationship to time, *α* called subdiffusion (1 > *α* > 0) or superdiffusion (*α* > 1) and *β* called measurement error.

**VSL-label**
|  | Ligand-free<br>(10,465) | Capsaicin<br>(8,026) | AMG9810<br>(10,578) |
| --- | --- | --- | --- |
| $D_x$<br>($\text{mrad}^2/\text{ms}$) | 4.03 | 5.88 | 5.68 |
| $\alpha$ | 1.37 | 1.51 | 1.38 |
| $\sigma^2$<br>($\text{mrad}^2$) | 0.59 | 0.86 | 0.53 |

**Table S1| Parameter of MSD for wild type of TRPV1**
|  | Ligand-free<br>(12,108) | Capsaicin<br>(10,660) | AMG9810<br>(11,090) |
| --- | --- | --- | --- |
| $D_x$<br>($\text{mrad}^2/\text{ms}$) | 4.30 | 4.39 | 4.61 |
| $\alpha$ | 1.49 | 1.57 | 1.68 |
| $\sigma^2$<br>($\text{mrad}^2$) | 0.75 | 0.83 | 0.85 |

**Table S2.** Parameter of MSD for each Lifetime group in OP-label with capsaicin. The MSD curve were fitted by the following function: *δ* ^2^(*t*) = *Dα tα* + 2*β* ^2^. *D*_*α*_ is the anomalous diffusion constant, a non-linear relationship to time, *α* called subdiffusion (1 > *α* > 0) or superdiffusion (*α* > 1) and *β* called measurement error.

|  | short | middle | long |
| --- | --- | --- | --- |
|  | LF:0.4-2.5 ms | LF:2.6-4.0 ms | LF:8.0-10 ms |
| $D_\chi$<br>(mrad<sup>2</sup>/ms) | 9.87 | 7.30 | 5.04 |
| $\alpha$ | 1.27 | 1.40 | 1.39 |
| $\sigma^2$<br>(mrad<sup>2</sup>) | 0.59 | 0.46 | 0.29 |

