## Supplementary figures and images for "Agonist and antagonist diverted twisting motions of single TRPV1 channel"

### Video1

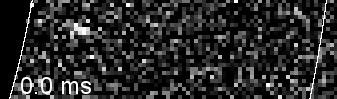

### Video2

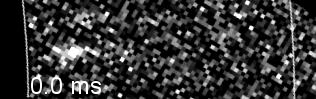
